# “Skull base meningiomas have a distinct immune landscape.”

**DOI:** 10.1101/525444

**Authors:** Zsolt Zador, Alexander P. Landry, Michael Balas, Michael D. Cusimano

## Abstract

Modulation of tumor microenvironment is an emerging frontier for new therapeutics. However in meningiomas, the most frequent adult brain tumor, the correlation of microenvironment with tumor phenotype is scarcely studied. We applied a variety of systems biology approaches to bulk tumor transcriptomics to explore the immune environments of both skull base and convexity (hemispheric) meningiomas. We hypothesized that the more benign biology of skull base meningiomas parallels the relative composition and activity of immune cells that oppose tumor growth and/or survival. We firstly applied gene co-expression networks to tumor bulk transcriptomics from 107 meningiomas (derived from 3 independent studies) and found immune processes to be the sole biological mechanism correlated with anatomical location while correcting for tumour grade. We then derived tumor immune cell fractions from bulk transcriptomics data and examined the immune cell-cytokine interactions using a network-based approach. We demonstrate that oncolytic M1 macrophages dominate skull base meningiomas while mast cells, known to play a role in oncogenesis, show greater activity in convexity tumors. Our results are the first to suggest the importance of tumor microenvironment in meningioma biology in the context of anatomic location and immune landscape. These findings may help better inform surgical decision making and yield location-specific therapies through modulation of immune microenvironment.

## Introduction

Meningiomas are the most common adult brain tumor and constitute approximately 30% of all intracranial neoplasms^1^. Surgery remains a key part of treatment for symptomatic or growing meningiomas and outcomes are largely determined by tumor biology^2,3^ and extent of resection^4^. Approximately 70% of meningiomas have benign characteristics and the average disease-free survival is 90% over 10 years^2^, but lifetime recurrence can be as high as 50%^5^. The extent of surgical excision is largely determined by technical feasibility which is a function of anatomical location of the tumor, adherence to adjacent tissue and the eloquent structures limiting the extent of removal^6^. Achieving complete excision of skull base meningiomas poses a particular challenge due to the proximity of neurovascular structures as well as the often narrow surgical corridors. Consequently, they may require longer operative times and can have lower success rate of complete excision with eradication of tumor origin. Therefore, understanding the biology skull base meningiomas is of particular interest given these patients are more likely to be left with residual disease.

Multiple studies show that skull base meningiomas are more likely to have benign biology whereas tumors with more aggressive behavior (atypical or malignant meningiomas) can constitute close to 30% of convexity/parafalcine tumors^7–9^. Recent studies have analyzed the genetic makeup of meningiomas and found recurrent mutations in the neurofibromatosis type 2 (NF2) gene and/or loss of chromosome 22 (NF2/chr22loss) to be more prevalent in the cerebral and cerebellar hemisphere^10^. The vast majority of non-NF2/chr22loss meningiomas are typically benign and tend to be located medially on the skull base. Smoothened, frizzled family receptor (SMO) mutations are also implicated in non-NF2/chr22loss and medial meningiomas through increased activation of the Hedgehog pathway^10^. Recurrent polymerase (RNA) II (DNA directed) polypeptide A (POLR2A) mutations have recently been found to classify a distinct subset of benign meningiomas with meningothelial histology and a preponderance to localize in the tuberculum sellae^11^. Analysis of methylation subtypes of meningiomas showed benign subgroups with longer disease-free survival had a greater proportion of skull base meningiomas. On the other hand, all tumors in the malignant methylation subtype were located exclusively on the convexity^8^.

Systems level analysis of gene expression data, which considers genes as units of a system rather than isolated entities, can identify key molecular processes which map to biological/clinical phenotypes^12,13^. This approach carries the distinct advantage of being unbiased in deriving meaning from high dimensional, biologically-modelled data. Gene co-expression networks are one example of such system level analysis, wherein similar genes are grouped into “modules” of similar function^14^. This technique has been successfully applied to explore complex phenotypes in Huntington’s disease^15^, peripheral nerve regeneration^16^ and weight gain^17^. It has also been used to identify relevant subgroups of meningioma^18^. In the current study we apply this technique along with another network-based approach to model the immune landscape of skull base and convexity meningiomas.

We hypothesize that different meningioma locations are associated with distinct biological mechanisms, likely related to immune cell composition and/or activity. To capture these geographical patterns in meningioma biology we analyzed the transcriptomics profiles of 107 meningiomas originating from either the skull base or convexity.

## Results

### Co-expression network and module-phenotype correlation

A gene co-expression network was constructed from 19,011 gene transcripts. Using an adaptive hierarchical clustering model^14^, we discovered fourteen gene modules (Figure 1A). We next compared the biology of both tumor locations by correlating phenotype with meta-gene expression levels. Three modules were found to be significantly different between hemispheric and skull base meningiomas (Mann Whitney p<0.05), only one of which annotated significantly (Bonferroni p<0.05) to biological processes in DAVID (Figure 1B,C). We finally confirmed module significance by regressing out WHO grade in a generalized linear model.

**Figure 1:**
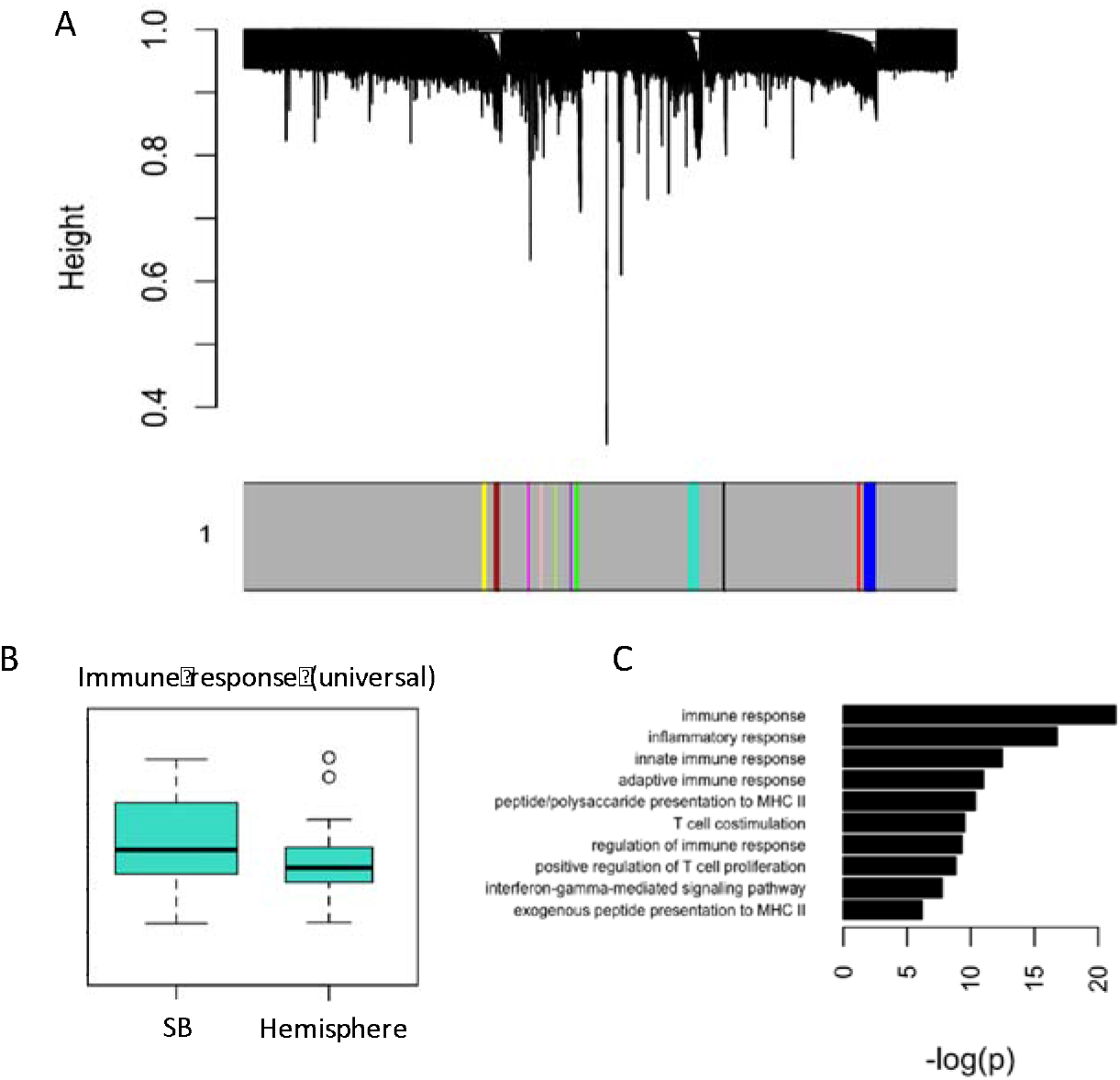
Gene co-expression network reveals immune function to correlate strongly with meningioma location. A: Gene dendrogram illustrating modules. The grey denotes genes which are not implicated with any modules. B: Boxplot depicting the correlation between location and meta-gene expression level of the only significant module with DAVID annotations (Mann Whitney p = 0.005). C: Gene ontology terms ranked by Bonferroni p-value.

### Immune composition of meningiomas by location

Using network analysis, we examined the biological activity of each immune cell type by correlating them with cytokine expression. On inspection of cytokine-cell networks there was a clear difference in the network configurations of convexity and skull base meningiomas (Figure 2). Notably, the convexity network yields two distinct cell-cytokine clusters (as well as an independent correlation between activated natural killer cells and resting mast cells). Monocytes are the sole cell type in one of the clusters, and are most correlated (ρ = 0.65) with CX3CR1, associated with leukocyte migration, cell adhesion, and monocyte differentiation/survival^19^ and CCR1, involved in monocyte and T cell chemotaxis^20^. In the other cluster, activated mast cells are most correlated (ρ = 0.81) with IL1RN, an antagonist of the interleukin-1 receptor^21^ and neutrophils are most correlated (ρ = 0.73) with CXCR2, a neutrophil-specific chemokine^22^. Both mast cells and neutrophils are connected to the pro-inflammatory cytokine IL1B^23^. In the skull base network, M0 macrophage associations are similar to the monocyte associations in the convexity model. Additionally, M1 macrophages are most correlated with CXCL10 (ρ = 0.74), another pro-inflammatory chemokine^24^.

**Figure 2:**
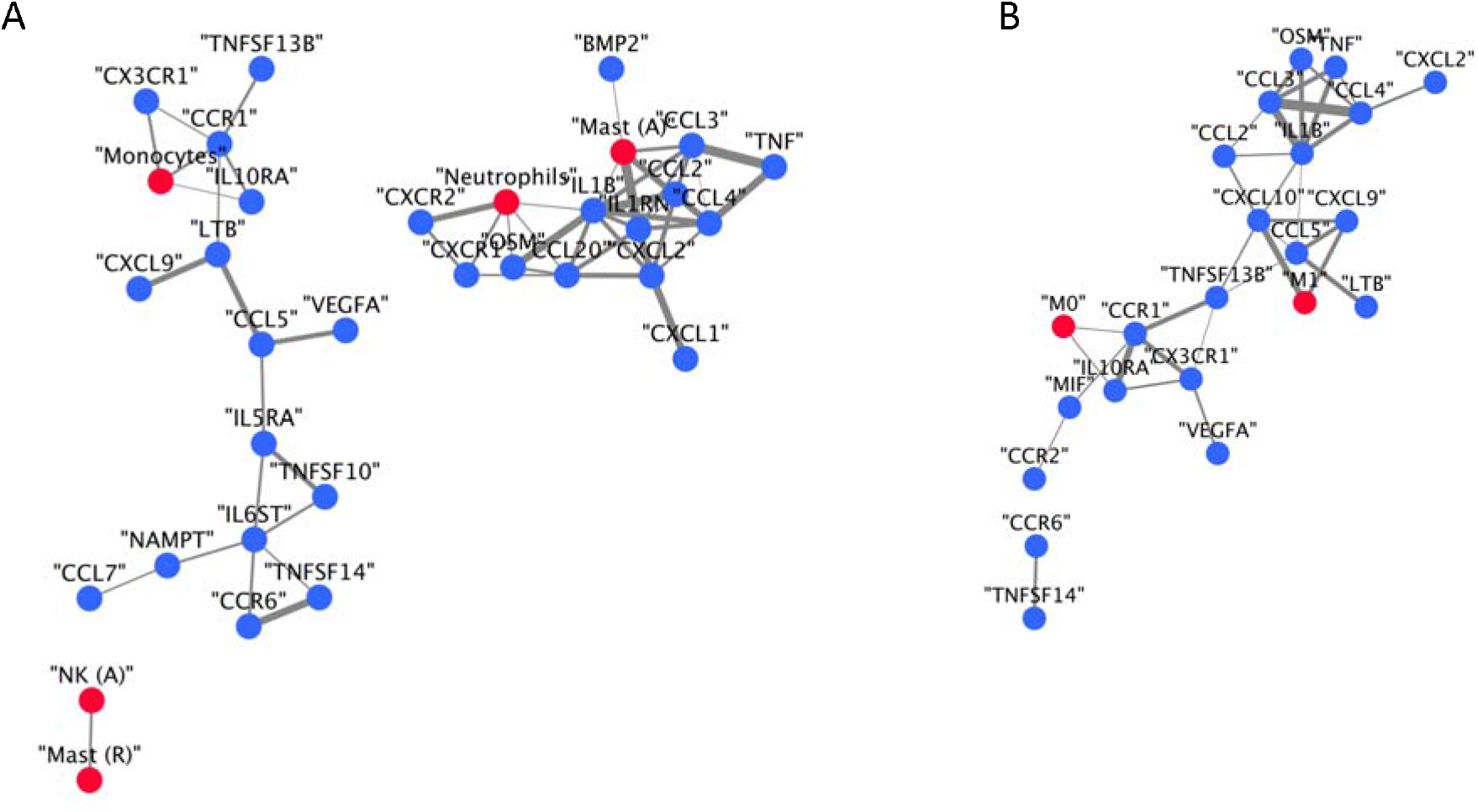
Network demonstrating the connectivity of immune cell fractions and cytokine transcriptomics in convexity (A) and skull base (B) meningiomas. Cytokines are represented in blue whereas immune cells are represented in red. Edge thickness is proportional to Pearson correlation. Neutrophils, monocytes, and mast cells are significantly correlated with cytokines in the hemispheric model, while M0 and M1 macrophages are significantly correlated with cytokines in the skull base model.

Given the relative complexity of these networks, we sought to analyze the relative “activity” of each immune cell by probing the distribution of their correlations with cytokine expression levels^25^. Analysis of cell “connectivity” (the area under this distribution) re-demonstrates the importance of mast cells and neutrophils in convexity meningiomas of and macrophages in skull base meningiomas (Figure 3).

**Figure 3:**
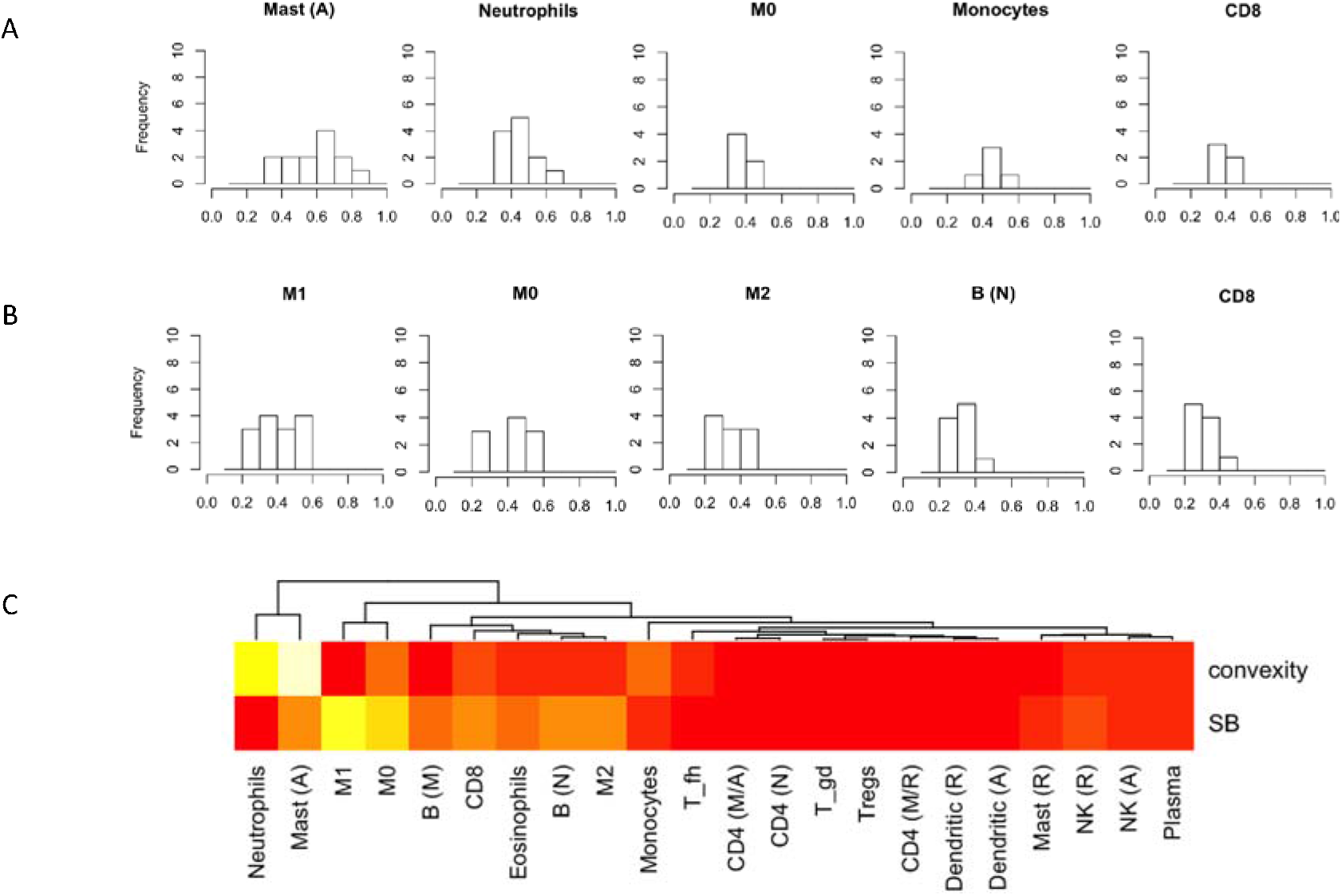
Immune cell connectivity of meningiomas by location. A-B: Histogram of Person’s correlations for the top 5 cell fraction, ranked by connectivity, of convexity (A) and skull base (B) meningiomas. C: Heatmap of cell connectivity values. White indicates high connectivity and red indicates low connectivity (range 0-7.8). Note the highly connected mast cells in convexity meningiomas and M1 macrophages in skull base (“SB”) meningiomas. A = activated, M = mature, N = naïve, fh = follicular helper, gd = gamma-delta, R = resting.

## Discussion

We present a genetic meta-analysis correlating transcriptomics profile with meningioma location. Using a systems biology approach, we demonstrate an upregulation of various immune processes in skull base meningiomas compared to convexity (hemispheric) meningiomas. We further investigated the immune landscape of meningiomas by combining leukocyte cell fraction with cytokine expression profile in a network-based analysis. M1 macrophages, known to have anti-tumor activity, were associated with the greatest cytokine connectivity in skull base meningiomas. This finding may explain their more benign biology compared to convexity meningiomas.

The more aggressive grade II and grade III meningiomas are less prevalent in the skull base compared to convexity^7,26^. When comparing to meningiomas of the convexity, only immune processes showed significant upregulation in meningiomas of the skull base which was maintained after correcting for histological grade. This finding parallels the concept that cellular environment modulates tumor biology. This process has been well demonstrated in a variety of pan-cancer^12,27^ and central nervous system tumour studies^28^, and has been suggested in meningiomas^29–31^. Multiparameter flow cytometry studies have demonstrated a heterogenous composition of immune cells in meningiomas^29,31^. More specifically, there was a variable degree of infiltrative macrophages with high phagocytic activity and, to a lesser extent, T-cells, Natural Killers, and few B-cells. Analysis of fresh meningioma tissue has demonstrated a large fraction of leukocyte infiltration consisting of macrophages with immunosuppressive activity^30^, which are more prevalent in grade II tumors. In our study, M1 macrophages were the most active cell fraction based on overall cytokine correlation. This cell fraction is also referred to as the “kill-type” macrophages and regarded as an inhibitor of tumor growth in multiple tumor types^32,33^. In convexity meningiomas, on the other hand, we identified activated mast cells as the most connected fraction. Tumor-associated mast cells may support the oncogenic environment by releasing pro-tumorigenic stimulants^34^ triggering angiogenesis, tumor cell proliferation/invasion, formation of lymphatic/blood vessel and facilitate the process of extravasation of cytokine-producing cells. Mast cells have been detected in 90% of meningioma tissue and has been also found to correlate with peritumoral edema^35^, a feature of aggressive biology in meningiomas.

Conventional analysis of transcriptomics data often relies on differential single-gene expression levels to filter out relevant patterns by comparing the “diseased” and “normal” tissue. These techniques may overlook relatively subtle effects across several highly-connected genes, which may nevertheless trigger a robust cellular mechanism. Gene expression networks are well suited for detecting small, additive biological signals that underlie relevant clinical phenotypes. In the first part of our analysis we used this well-established technique^14^. We found that immune processes correlate with location of meningiomas and investigated this finding further using an integrated, network-based approach. In this second technique, rather than using the conventional approach of considering cell fractions as a reflection of immune landscape we included correlations of cytokine expression patterns with cell fractions. We used the sum of cell-cytokine correlations as a measure of immune cell “activity”, which is inspired by existing network based approaches^25^. Although further verification of this approach is needed we highlighted distinct cell fractions that have previously been shown pro- and antitumor activities and parallel biology of skull base meningiomas versus those of the convexity.

Limitations of our study are that it’s based purely on computational data, though it’s nevertheless in alignment with prior knowledge on tumor immunology and the biology of convexity vs skull base meningiomas. Our assessment of cytokine activity is on the transcriptomics level as no secretory data was available, yet this approach still provides an indirect assessment of how “active” an immune cell type is in terms of cytokine synthesis. Finally, the proportion of grade II meningiomas is relatively low in our study (5.3%) compared to the prevalence in the population (20-30%). To address this, we have corrected for grade when correlating module gene expression with location. Importantly, we have been able to identify a robust, recurrent signal despite considerable data heterogeneity.

## Conclusions

Our study is the first to computationally estimate immune cell fractions in location-specific meningioma tissue from bulk transcriptomics. We demonstrate distinct immune compositions between hemispheric and skull base meningiomas using a network-based approach that considers cell connectivity with cytokine transcriptomics. Macrophages with cytotoxic activity are more dominant in skull base meningiomas, in keeping with a more benign biology. These findings give further insight into the immune microenvironment of meningiomas, and have implications on future strategies of immune modulation for this challenging disease.

## Methods

### Data Preparation

All data was collected from the Gene Expression Omnibus (GEO), a public repository of high-throughput functional genomic data sets^36^. We used studies containing details on meningioma location (skull base and convexity) and WHO grade with corresponding gene expression data^11,37^, providing us with a cumulative sample size of 107 meningiomas (Table 1). Summary statistics for each study are presented in Table 1. The average age of our cohort is 54.6 (missing age values for GSE84263 were imputed for each), and percentage proportion of males to females is 22.4% to 77.6%, respectively.

**Table 1.**
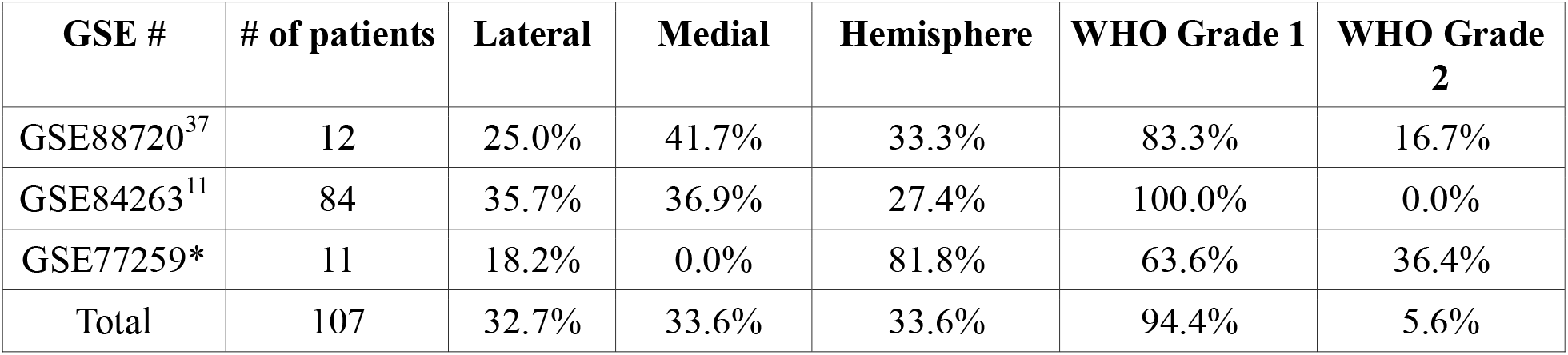
Table summary of study population.*No associated publication

### Pre-processing of transcriptomics data

For each study, the microarray data was backgrounded corrected, quantile normalized, and log-2 transformed using the *Affy*^38^ and *Limma*^39^ R packages for Affymetrix and Illumina platforms, respectively. After removing genes that were not common across these studies we were left with 19,011 genes. The 3 studies were merged, scaled to a global mean and standard deviation of 0 and 1, respectively^40^, and batch-corrected using *ComBat*, a well-established empirical Bayes approach^41^. The resultant data matrix was used during all subsequent analysis.

### Co-Expression Network Analysis

We performed WGCNA of gene expression data using R (version 3.5.1) to construct a co-expression network and identify biological modules which map to meningioma location. Pairwise gene correlations were soft-thresholded with an exponent of 20 to approximate scale-free topology, which was ultimately transformed into a biologically-inspired “Topological Overlap Matrix” (TOM), which measures pairwise gene similarity in terms of shared topology within the full network^14^. Highly similar genes are then grouped into an adaptive hierarchical clustering tree (dendrogram), yielding “modules” with a minimum size of 30 genes. The gene expression profile of each module is represented by a meta-gene computed as its first principal component, an established method.

### Module-Based Qualitative Analysis

Genes in each of the identified modules were annotated by the Database for Annotation, Visualization and Integrated Discovery (DAVID version 6.8)^42^.

### Deconvolution of tumor bulk expression signal

We used CIBERSORT, a well-established technique to estimate immune cell fractions from our gene expression data. Briefly, the method compares mixed-cell population transcriptomics with an established signature matrix using support vector regression in order to estimate the relative prevalence of 22 immune cells. This technique is described in detail in ref 32^43^.

### Network analysis of immune cell fractions and expression of cytokines

The complex relationships between immune cells and cytokine expression levels were visualized using network analysis, which demonstrates associations that are otherwise difficult to appreciate. Such network-based approaches have revolutionized research into phenom-genotype similarities and elucidated the genetic basis of diseases^46,47^, predicted pathways of disease progression^25^, and identified novel drug targets^44^. Nodes represent immune cells (CIBERSORT output) and list of 35 known cytokines derived from the literature^27,48^ (Supplementary data). Edges represent Pearson correlations, wherein a significance cut-off of <0.05 and effect size cutoff of >0.6 are used. We define “connectivity” as the biological activity of each immune cell (i.e. its overall expression of cytokines), computed as the sum of all significant (p<0.05) Pearson correlations between a cell type and the expression levels of 35 pre-determined cytokines (Supplementary data). Cell types were ranked based on this metric.

## Acknowledgements

The authors would like to thank Dr. Claudia Dos Santos and Dr. Christopher Walsh for their assistance with analytic techniques.

## Author Contributions

ZZ: study design, manuscript preparation

APL: data analysis, manuscript revisions

MB: data analysis, manuscript revisions

MDC: manuscript revisions

## Availability of Materials and Data

All data referenced in this study is publicly available through the open repository Gene Expression Omnibus (https://www.ncbi.nlm.nih.gov/geo/).

