## Supplementary table for "“Skull base meningiomas have a distinct immune landscape.”"

Cytokines:

CCL2, CCL3, CCL4, CXCL1, CXCL8, IFNG, IL10, IL12A, IL12B, IL1B, IL1R2, IL1RN, IL4, IL6, IL6R, TNF, TNFRSF1A, TNFRSF1B, BTK, CD244, CD274, CSF1, CSF2, EGF, GZMB, HAVCR2, IFNG, IL13, IL1A, LAG3, MMP9, PDCD1, PLAU, THBD, VEGFA
